# Analysis of Preplatelets and Their Barbell Platelet Derivatives by Imaging Flow Cytometry

**DOI:** 10.1101/2021.07.30.454421

**Authors:** Sam Kemble, Amanda Dalby, Gillian C Lowe, Phillip LR Nicolson, Steve P Watson, Yotis Senis, Steven G. Thomas, Paul Harrison

## Abstract

Circulating large “preplatelets” undergo fission via barbell platelet intermediates into two smaller, mature platelets. In this study, we determine whether preplatelets and/or barbells are equivalent to reticulated/immature platelets by using ImageStream flow cytometry (ISFC) and super-resolution microscopy. Immature platelets, preplatelets and barbells were quantified in healthy and thrombocytopenic mice, healthy human volunteers, and patients with immune thrombocytopenia (ITP) or undergoing chemotherapy. Preplatelets and barbells were 1.9%+0.18/1.7%+0.48 (n=6) and 3.3%+1.6/0.5%+0.27 (n=12) of total platelet counts in murine and human whole blood, respectively. Both preplatelets and barbells exhibited high expression of HLA-I with high thiazole orange and mitotracker fluorescence. Tracking dye experiments confirmed that preplatelets transform into barbells and undergo fission *ex vivo* to increase platelet counts, with dependence upon the cytoskeleton and normal mitochondrial respiration. Samples from antibody-induced thrombocytopenia in mice and patients with ITP had increased levels of both preplatelets and barbells correlating with immature platelet levels. Furthermore, barbells were absent post-chemotherapy in patients. In mice, *in vivo* biotinylation confirmed that barbells, but not all large platelets, were immature. This study demonstrates that a subpopulation of large platelets are immature preplatelets that can transform into barbells and undergo fission during maturation.

## Introduction

Platelets are anucleated discoid blood cells that originate from bone marrow megakaryocytes(MK).^1^ As human platelets exhibit a short lifespan of ∼10 days, over 100 billion new platelets are required per day to sustain the platelet count(150–400 × 10^9^/ L).^2^ The youngest platelets are termed reticulated platelets(RP),^3^ and were first described in acute blood loss.^4^ RP have been traditionally measured by flow cytometry using nucleic acid dyes(e.g. thiazole orange). ^5^ Biotinylation studies performed in mice have also confirmed that they are immature platelets.^3^ RP levels can distinguish between thrombocytopenic conditions due to enhanced peripheral destruction or lack of production.^6,7^ Commercial methods for quantification include the immature platelet fraction(IPF),^8,9^ which although standardised tend to overestimate the number in macrothrombocytopenia and bone marrow suppression/failure due to non-specific labelling of nucleotides.^10,11^ Recent efforts have tried to overcome such limitations identifying HLA-I(MHC-I) as a potential new marker of platelet immaturity.^12^

During maturation, MKs form long proplatelet extensions, which can subsequently release platelets.^13-15^ These proplatelets are released into the bone marrow sinusoids, with platelet maturation continuing within the bloodstream^15-18^ and the lungs.^19,20^ Furthermore, proplatelet-like barbell structures (consisting of figure of eight microtubules) undergo fission *in vitro* to form progeny, thus increasing the count.^21^ This led to the discovery of a new terminal stage of maturation which takes place in the bloodstream through a platelet intermediate the “preplatelet”.^17^ Balduini has recently highlighted that similar structures and platelet division were originally described by Perroncito back in 1921.^22^ Circulating preplatelets are large platelet precursors(3-10 μm) which divide by converting into barbells before undergoing fission into mature platelets (<3 μm)^17,21^ These structures have been identified and quantified in healthy human platelet-rich plasma (PRP) using laser scanning cytometry (3.6% and 0.05% respectively).^17,23^ However, barbells have been rarely reported probably because both IPF and blood film analysis tend to use EDTA anticoagulated blood, causing swelling and irreversible conversion of barbells back into preplatelets.^23^

Although the current dogma is that preplatelets and barbells represent intermediate structures that undergo fission, it is still unknown whether this exclusively occurs within the immature platelet population and there remains a need for a robust method to study the two species. Therefore, we applied ImageStream flow cytometry(ISFC) to detect and quantify both preplatelets and barbells directly in whole blood. ISFC allows full preplatelet and barbell characterisation as it combines the speed, sensitivity and phenotyping abilities of conventional flow cytometry(CFC) with single cell morphological and content resolution. By designing a new ISFC method to accurately discriminate preplatelets and barbells, for the first time we have quantified and characterised these structures in health and thrombocytopenic human and mouse blood and demonstrated their equivalence to immature platelets. We also confirmed that these structures can undergo terminal maturation via fission and propose morphometric quantification of preplatelet-derived barbells could be an additional tool to platelet counts and IPF for potentially diagnosing and managing thrombocytopenia.

## Materials and Methods

### Patient Recruitment

This study was approved by the NHS Research and Ethics Committee(NHS REC; 15/WM/0465) and by the University Hospitals Birmingham NHS Foundation Trust(RRK 5677). Healthy controls, patients with Immune Thrombocytopenia(ITP), and those with haematological malignancies undergoing chemotherapy-induced bone marrow ablation were recruited. Blood samples were anticoagulated in EDTA, trisodium citrate and hirudin BD Vacutainers®(Fisher Scientific and Roche respectively). See supplementary methods for patient demographics.

### Mice

Wildtype(WT) C57BL/6 mice were of mixed gender between 8-12 weeks old. All animal procedures were undertaken with UK Home Office approval under a project licence(P46252127).

### Antibodies and Probes

Whole blood or washed platelets were labelled with anti-human BV421 CD62p (P-selectin; 1/100, AK4, BioLegend, 304926), BV421 CD42b(1/300, HIP1, BioLegend, 303930), FITC CD61(β3; 1/25, RUU-PL7F12, BD, 348093), Thiazole Orange(200ng/mL; Sigma, 390062), PE CD42b(GPIbα 1/50, HIP1, BD, 555437), PE CD62p(1/100, AK4, BioLegend, 304906) AF599 mitotracker Ros CMX(mitochondrial dye 5μM, Cell Signalling, 9082), APC CD62p(1/100, AK4, BioLegend, 304910) and/or APC HLA I(1/100, W6/32, BioLegend, 311410); or anti-mouse BV421-CD62p(1/100, VI P-44, BioLegend, 304926), FITC-CD41a(1/100, MWReg30, BioLegend, 133904), FITC-conjugated streptavidin(1/10, BioLegend, 405201), BD Retic-count (BD, 349204) and/or APC-CD41(1/100, MWReg30, ThermoFisher, 17-0411-82). A Polyclonal rat anti-mouse GPIb antibody(Emfret, R300) was used to induce thrombocytopenia in mice.

Isotypes BV421 mouse IgG1(1/100, MOPC-21, BioLegend, 400157), BV421 rat IgG1(1/100, A110-1, BD; 562604), PE or APC mouse IgG1(1/100, MOPC-21, BioLegend, 400112 and 400120 respectively) were used to quantify CD62p exposure. Fluorescence minus one was used for all other markers. AF674 SiR tubulin probe(4μM SpiroChrome, #CY-SC002) was used for ISFC tubulin labelling.

### Haematology Analysers

Human and mouse samples were analysed on the XN1000(Sysmex UK) and ABX Pentra 60(Horiba) haematology analysers respectively. Platelet counts, IPF and mean platelet volume (MPV) were recorded.

### Blood Smears

Human whole blood was stained with modified Giemsa stain(see manufacturer’s guidelines; Gentaur).

### Washing Platelets

Platelets were washed as previously described.^24^ Following washing steps, platelets were resuspended and diluted in serum-free M199 media to 100 × 10^6^/mL.

### Platelet Incubations

For all incubations, platelets were incubated in M199 media(ThermoFisher, 31150022) and maintained in suspension using a microplate mixer(300 rpm; StarLab N2400-8040). Incubations were either conducted on the bench(21°C) or within a cell incubator(37°C). For whole blood experiments, 1mL of blood was incubated in a 5mL polypropylene tube for 0, 0.25, 1.5 or 3h. For washed platelets, 100 × 10^6^/mL platelets were incubated for 0, 1.5, 3, 6 or 24h.

To inhibit platelet aggregation, whole blood was incubated with 10μM eptifibatide (Sigma, SML1042) and incubated for 3h at 21°C. For disruption of cytoskeletal signalling washed platelets were incubated with nocodazole(10μM; Sigma, M1404) or cytochalasin D(1 or 0.1μM; Sigma, C6762). To inhibit mitochondrial respiration washed platelets were treated with rotenone(3μg/mL; Sigma, R8875). For dual cytosolic labelling washed platelets were incubated with CellTrace™ CFSE (green) or red(0.2μg/mL and 1μg/mL; ThermoFisher, C34570 and C34572, respectively) for 30 minutes and centrifuged at 1000g for 10 minutes with 1μM prostacyclin I_2_(PGI_2_; Cayman, 61849-14-7). Green and red labelled platelets were resuspended in serum free M199 media at 100 × 10^6^/mL, mixed and incubated for 3h. Similar experimental conditions and methodology were used to track preplatelet maturation using labelling with CellTrace™ yellow(2μg/mL; ThermoFisher, C34573).

### Thrombocytopenia in mice

To induce thrombocytopenia, C57BL/6 mice received an intraperitoneal injection of polyclonal rat anti-mouse GPIbα antibody (final concentration of 1.5 μg/g). Platelet counts were measured from tail bleeds on day 0 (prior to challenge), 1, 5 and 8. For preplatelet and barbell ISFC analysis, mice were culled by terminal bleed (in 1/10 volume 4% trisodium citrate) on day 5.

### In vivo platelet biotinylation

Wild type C57BL/6J mice received two intravenous (IV) injections of NHS-biotin(4 mg/mL (Sigma, H1759), in saline) 30 minutes apart to ensure the circulating and splenic pools of platelets were labelled. A tail bleed was performed at 1h and 24h post second IV injection to measure percentage biotin positive platelets. At 24h mice were also culled via terminal bleed and whole blood was anticoagulated in 1/10 volume of 4% trisodium citrate.

### Flow cytometry

Reticulated and biotin positive platelets were measured using the Accuri™ C6 (see supplementary methods).

### Imaging Flow cytometry

ISFC was performed using the ImageStream^X^ MKII single camera system with brightfield, side scattered light and lasers 405 (60mW), 488 (80mW) and 642nM (100mW). Images were captured using an x60 objective lens (pixel area of 0.1μm^2^, maximum field of view of 128 μm). Acquisitions consisted of 10,000 platelet images (anti-human FITC CD61(Figure 2A & 3-5) or BV421 CD42b(Figure 2B only) or anti-mouse FITC or APC CD41 fluorescence). For human preplatelet and barbell measurements(Figure 2A & 3-5), antibody panels consisted of BV421 CD62p, Channel 1 (CH01); FITC CD61, CH02; AF599 mitotracker Ros CMX, CH04 and brightfield, CH04 or CH06; For tubulin analyses, BV421 CD62p in, CH01; FITC CD61, CH02 and AF674 SiR-Tubulin, CH05. For immature platelet markers, BV421 CD42b, CH01; TO, CH02; Mitotracker Ros CMX, CH03; PE CD62p CH04; APC HLA-I, CH05 and SSC, CH06. For mice, preplatelet and barbell analyses BV421 CD62p, CH01; FITC CD41, CH02 and brightfield, CH04 (Figure 6A-D); or for biotinylation BV421 CD62p, CH01; FITC streptavidin, CH02; brightfield, CH04 and APC CD41, CH05 (Figure 6E-F).

Captured images were analysed and optimised(fluorescent RMS gradient >20) on IDEAS(Image Data Exploration and Analysis Software). To determine barbell gating, truth populations, single circular and barbell platelets were manually selected and individually combined with pixel masks(morphology (includes all pixels from within the outermost contour), skeleton thin (a 1-pixel wide skeletal line within an object) and erode (removes the outermost pixels from the default mask)) to generate relative difference(RD) statistical feature tables. Features were chosen based on RD scores and visualised to determine optimal feature parameters to discriminate barbells from all other platelets. Preplatelets and barbells were determined by features include intensity (the sum of pixel intensities within a mask), aspect ratio (ratio of the minor axis divided by the major axis; 1=circular), area (size of the mask in μm^2^), diameter (diameter of a circle which is the same area as the object), compactness (density of pixel intensities within the object), symmetry2 (tendency of an object to have two lobes with a single axis of elongation), height (based on a bounding rectangle where the longer side is the height) or minor and major axis intensity(narrowest or widest region of an object).

For preplatelet and barbell gating, all analyses were performed on CD62p negative platelets to exclude microaggregate contaminants. For preplatelet quantification a CD61, CD41 or CD42b morphology pixel mask was designed to generate features aspect ratio and area of platelet images. Preplatelets were determined by an aspect ratio score of 0.8 – 1. From this an erode+4-pixel mask was designed to determine preplatelets with a diameter ≥3μm. For barbell quantification resting elongated platelets were discriminated from circular/discoid platelets using compactness and height*symmetry2 feature measurements of the CD61 morphology mask. From this, a skeleton thin mask was designed to depict a single pixel thick cytoskeletal line of the barbell structure and barbells were determine by a low minor axis intensity of CD61 morphology and high area*minor axis intensity of the CD61 skeleton thin (the barbell gate). See supplementary Figure 3 for visualisation and description.

As no barbell platelets were present at 21°C, characterisation of barbell-shaped microaggregates was completed in 21°C citrate blood (using EDTA blood as a negative control) using a similar ISFC methodology to the barbell analysis where microaggregates were discriminated from elongated platelets by a high symmetry2 and fluorescence intensity of a CD61 morphology mask.

### Immunofluorescence microscopy See supplementary methods Statistical Analysis

All data analysis was performed using GraphPad Prism v8.4.3GraphPad Software, California, USA). Statistical analyses are described within the figure legends.

## Results

### Preplatelets and barbells are immature reticulated platelets in human blood

In this study, a novel strategy using ISFC was developed to accurately detect and quantify preplatelets and barbells in whole blood, as described in the Supplementary Figures 1-3(details on pre-analytical variables: labelling, detection and gating). Preplatelets are detected as rounded structures which can be distinguished and resolved from platelets by their size(>3μm in diameter), while barbells have a unique “figure of eight” appearance which distinguishes them from platelet doublets/microaggregates that are CD62p positive and composed of 2 individual platelets(Supplementary figure 1-2). Figure 1A(i-iii) shows several examples of labelling of the marginal tubulin band and CD61(integrin subunit β3) in platelets, preplatelets and barbells.

**Figure 1:**
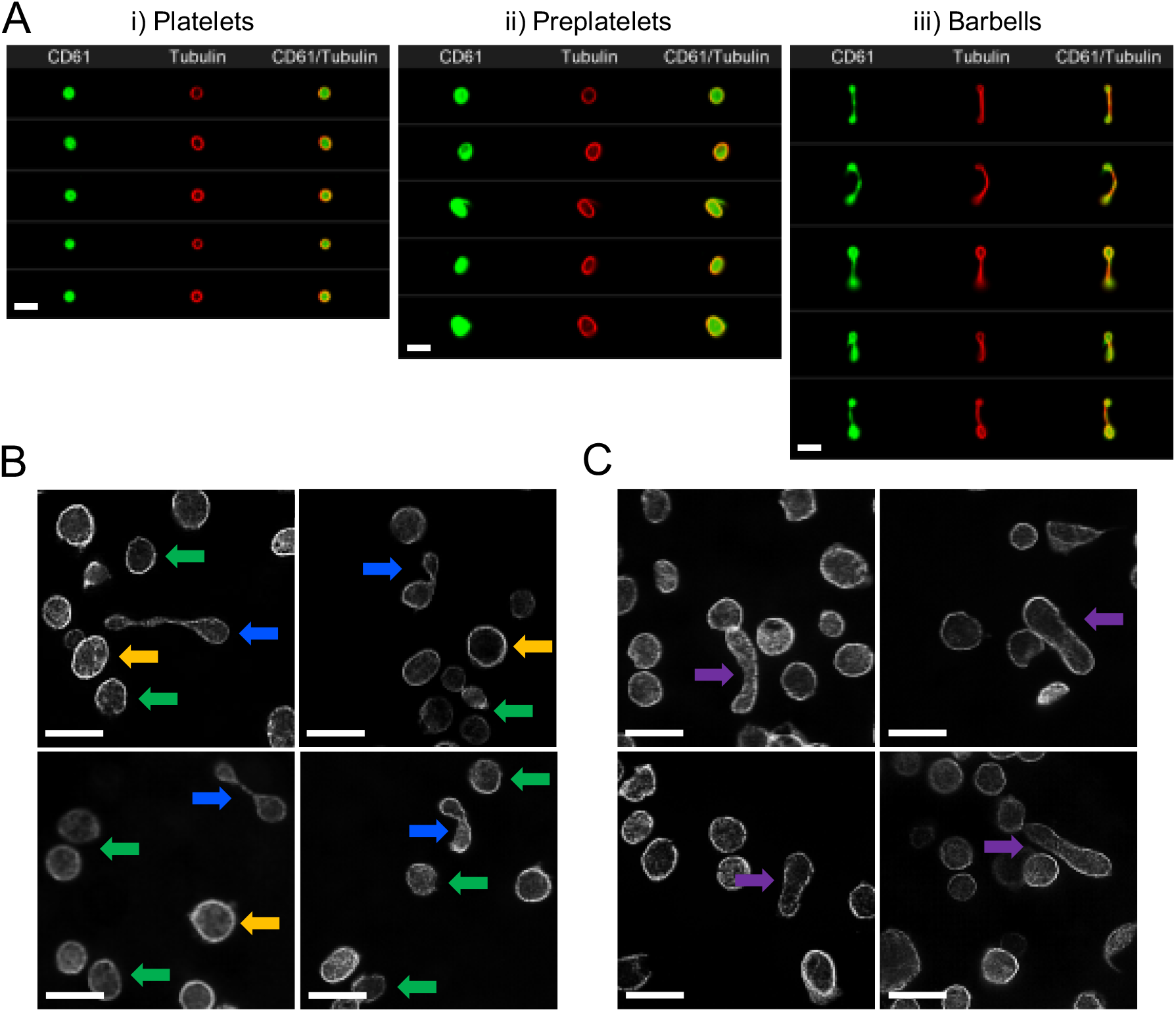
Imaging of Platelets, preplatelets and barbell platelets in normal human blood. A) Healthy control trisodium citrate blood was labelled immediately after phlebotomy with FITC anti-CD61 and AF674 SiR tubulin (4μM) for 30 minutes at 37°C and 10,000 CD61 positive platelet images were acquired using ISFC. SiR tubulin labelling clearly depicts the marginal band of (A) platelets, (B) preplatelets and (C) barbell platelets discriminated by ISFC (x60). Images are representative of a single experiment (n=5). Scale bar = 7μm. B) PRP was separated form citrated whole blood immediately after phlebotomy at 37°C and was labelled for α-tubulin and imaged using super-resolution SIM microscopy (x100 magnification, n=3). Circular platelets (≤ 3μm; green arrows) and preplatelets (≥ 3μm; yellow arrows) are shown along with barbell platelets (blue arrows). C) Circular platelets are shown along with Intermediate elongated preplatelets (purple arrows). Scale bars= 5μm

**Figure 2:**
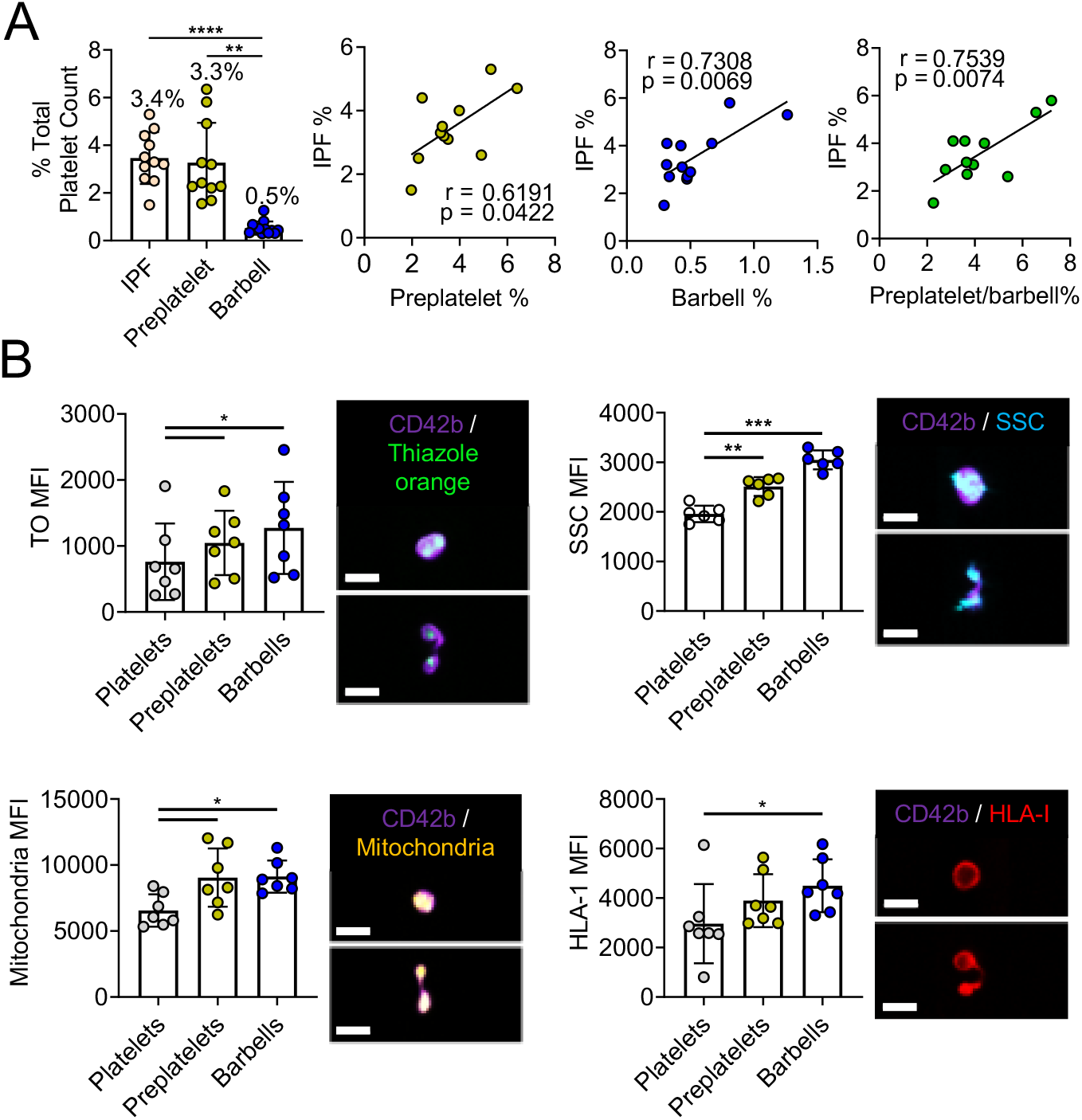
Quantification of reticulated platelets, preplatelets and barbells in human whole blood. (A) Healthy control trisodium citrate whole blood was labelled immediately after phlebotomy with anti-CD61 FITC and CD62p BV421 for 15 min at 37°C and percentages of immature platelets (IPF) was measured by Sysmex XN1000 and, preplatelets and barbells were quantified by ISFC and correlated with IPF (n=12).). (B) Platelets were then labelled with anti-CD42b BV421 and thiazole orange (TO), mitotracker AF599 or anti-HLA APC (n=5; scale bar 7μm) and granularity was determined by side scattered light SSC. Representative images of preplatelets and barbells with each label are shown (B). MFI was normalised to cellular perimeter. IPF: immature platelet fraction. (A & B) One-way anova with Dunnett’s multiple comparisons test; (B) Pearson’s correlation coefficient. Sig. * <0.05, ** <0.01, *** <0.001, ****<0.0001. +/-1 SD.

**Figure 3:**
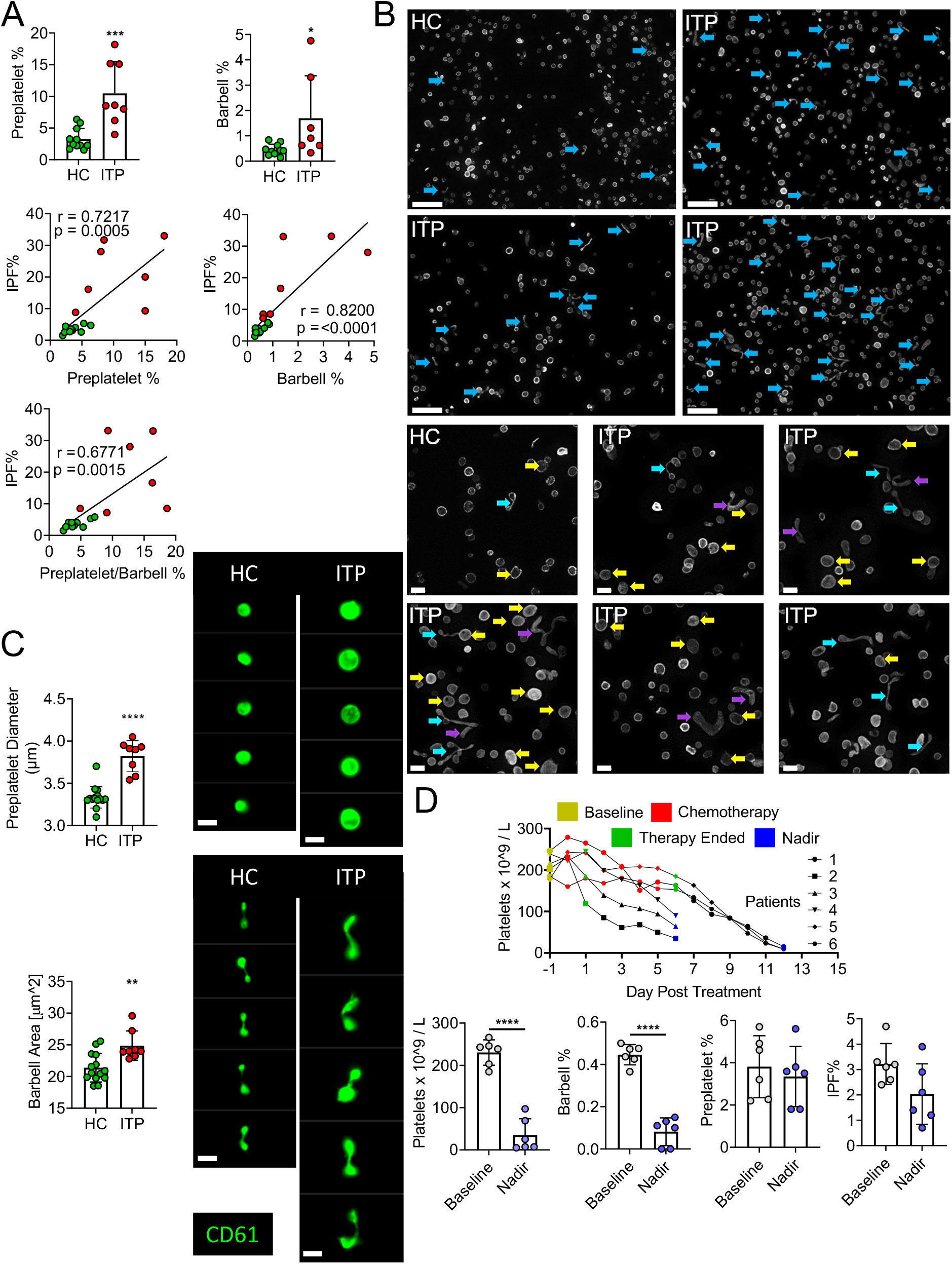
Quantifying reticulated platelets, preplatelets and barbell proplatelets in acquired thrombocytopenia. All measurements were performed in trisodium citrate anticoagulated whole blood. (A) Healthy control (HC; green dots: n=12) and immune thrombocytopenia (ITP; red dots: n=7) blood was incubated at 37°C for 1.5h and platelet count, MPV and IPF was measured by the XN1000 haematology analyser (Sysmex) and preplatelets and barbells by ISFC with correlations between % reticulated platelets (IPF%) and preplatelets and/or barbells. (B) α-tubulin immunofluorescence imaging of control and ITP PRP displaying preplatelets (yellow arrows), barbells (blue arrows) and elongated preplatelets (purple arrows; n=3; Leica DM6000 widefield microscope, x60 lens, scale bar 5μm). (C) Mean diameter of preplatelets and area of barbell platelets measured by ISFC comparing healthy controls (n=15) with ITP (n= 8; scale bar = 7μm). (D) Blood samples were taken from patients with high grade lymphoma prior to chemotherapy (baseline; day -1), 5-7 days post stem cell autograft (nadir). Platelet counts and IPF were measured by the XN 1000 analyser and preplatelets and barbells by ISFC (n=6). (A, B & D) Unpaired t-test; (A) Pearson’s correlation coefficient. Sig. * <0.05, ** <0.01, *** <0.001; **** <0.0001. +/- 1 SD.

**Figure 4:**
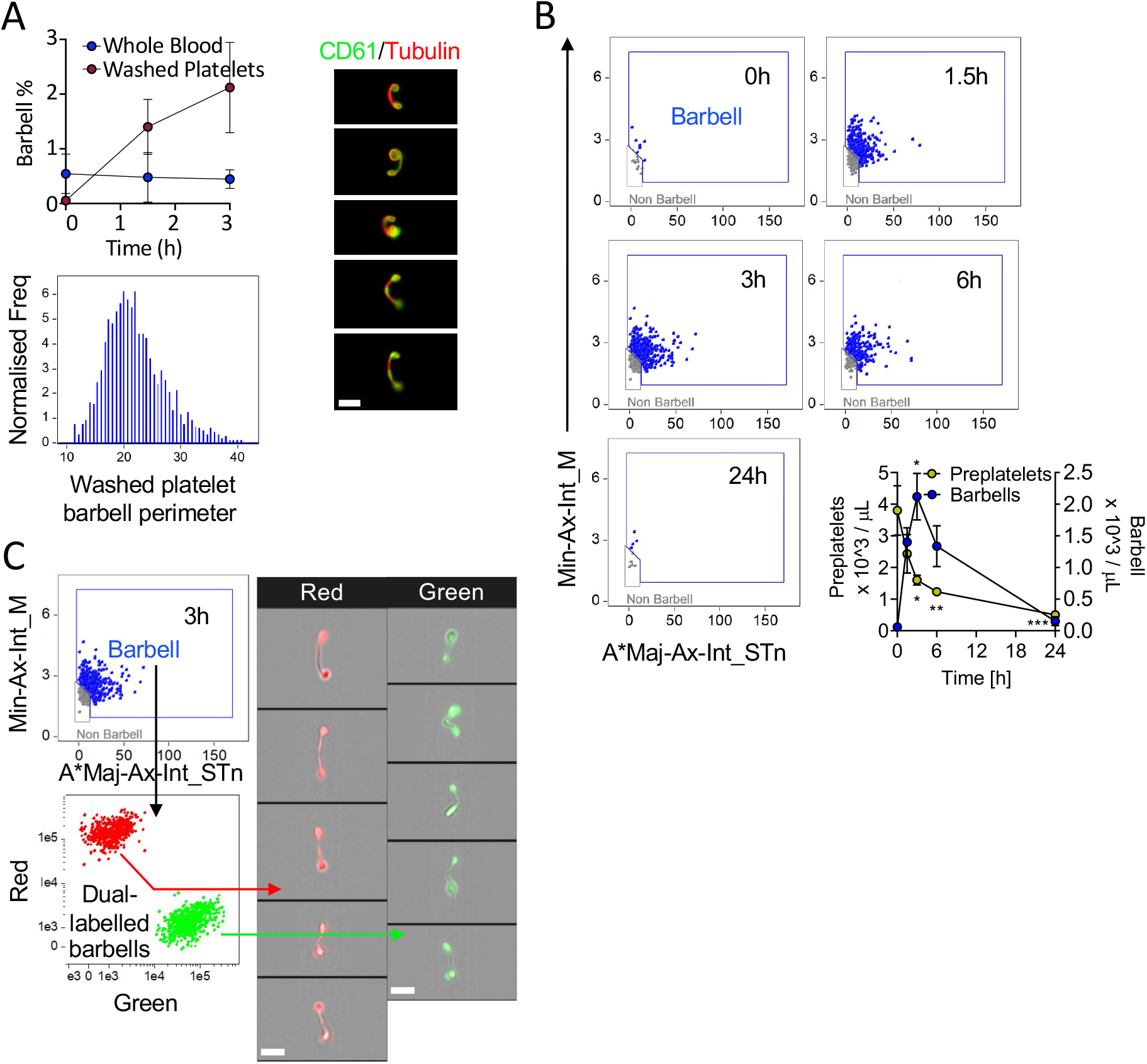
Kinetics of preplatelet maturation. (A) Whole blood and washed platelets anticoagulated with trisodium citrate was incubated for 3h at 37°C to determine change in barbell formation quantified by ISFC at 0, 1.5 & 3h and labelled with FITC anti-CD61 and AF674 SiR tubulin (4μM; n=6). Washed platelet barbell perimeter measured by ISFC (n=6). SiR tubulin live cell labelling of washed platelet barbells (n=3, scale bar 7μm). (B & C) Experiments were conducted with washed platelets from human control citrate blood incubated in M199 media at 37°C for a maximum of 24h. (B) Barbell platelet formation at 0, 1.5, 3, 6 and 24h time points (visualised using the image flow cytometry barbell gate described in Supplementary Figure 3). Quantification of preplatelets and barbells at 0, 1.5, 3, 6 and 24h time points (n=10). (C) Prior to incubating, washed platelets were labelled with either CellTrace™ green (0.2μg/mL) or red (1μg/mL) cytosolic dyes, mixed and incubated for 3h to demonstrate barbells originate from single platelets (n=3, imaged by ISFC, x60 lens, scale bar = 7μm). A*Maj-Ax-Int: Area*Major Axis Intensity; Min-Ax-Int: Minor Axis Intensity. (B) Two-way anova with Bonferroni multiple comparisons test. Sig *<0.05, **<0.01, ***0. +/- 1 SD.

**Figure 5:**
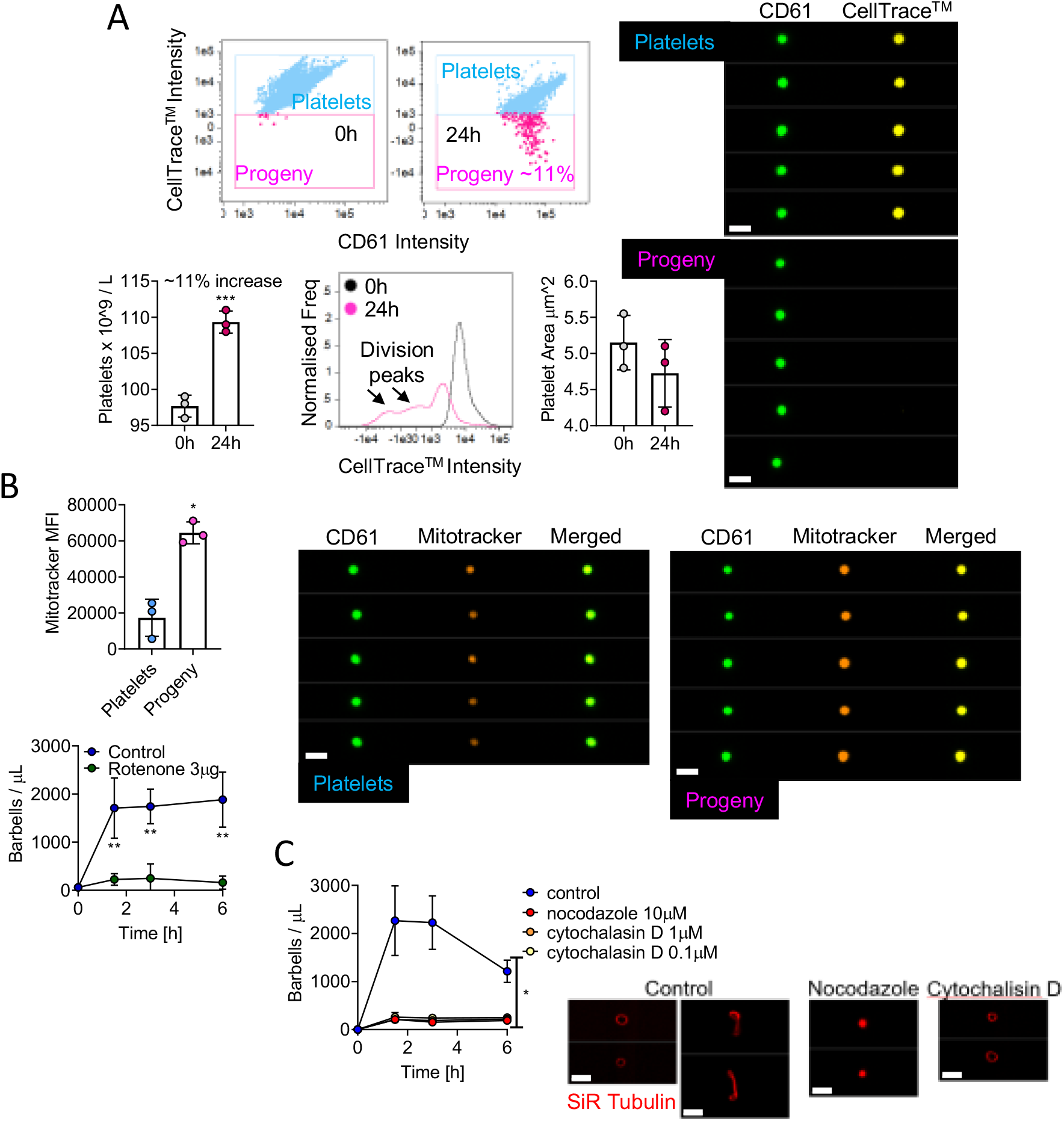
Mapping preplatelet maturation *in vitro*. Washed platelets were labelled with 2μg/mL of a CellTrace™ yellow cytosolic dye and incubated in serum-free M199 media for 24h. (A) Scatter plots demonstrate the appearance of a discrete population of platelets termed “Progeny” which display a decrease in CellTrace™ yellow mean fluorescence intensity (MFI; images depicted by CD61 and CellTrace™ fluorescence using ISFC, x60 lens, scale bars = 7μm). Percentage increase in platelet count, CellTrace™ fluorescence profile and platelet size at 0 and 24h are also demonstrated. (B) AF599 Mitotracker Ros CMX MFI of platelets and platelet progeny with representative images (scale bar 7μm) and the effect of rotenone (3μM) on barbell formation when incubating washed platelets for 6h. (C) Washed platelets were incubated for 6h with nocodazole (10μM) or cytochalasin D (1 or 0.1μM) and barbells were quantified by ISFC. To show the effect of either cytoskeletal drug, marginal band morphology was depicted using AF674 SiR tubulin live cell labelling (n=3). (A) paired t-test; (B) Mann-Whitney U test and Wilcoxon test; (C) two-way anova. Sig. ***=<0.001; **<0.01 & *<0.05. +/- 1 SD.

**Figure 6:**
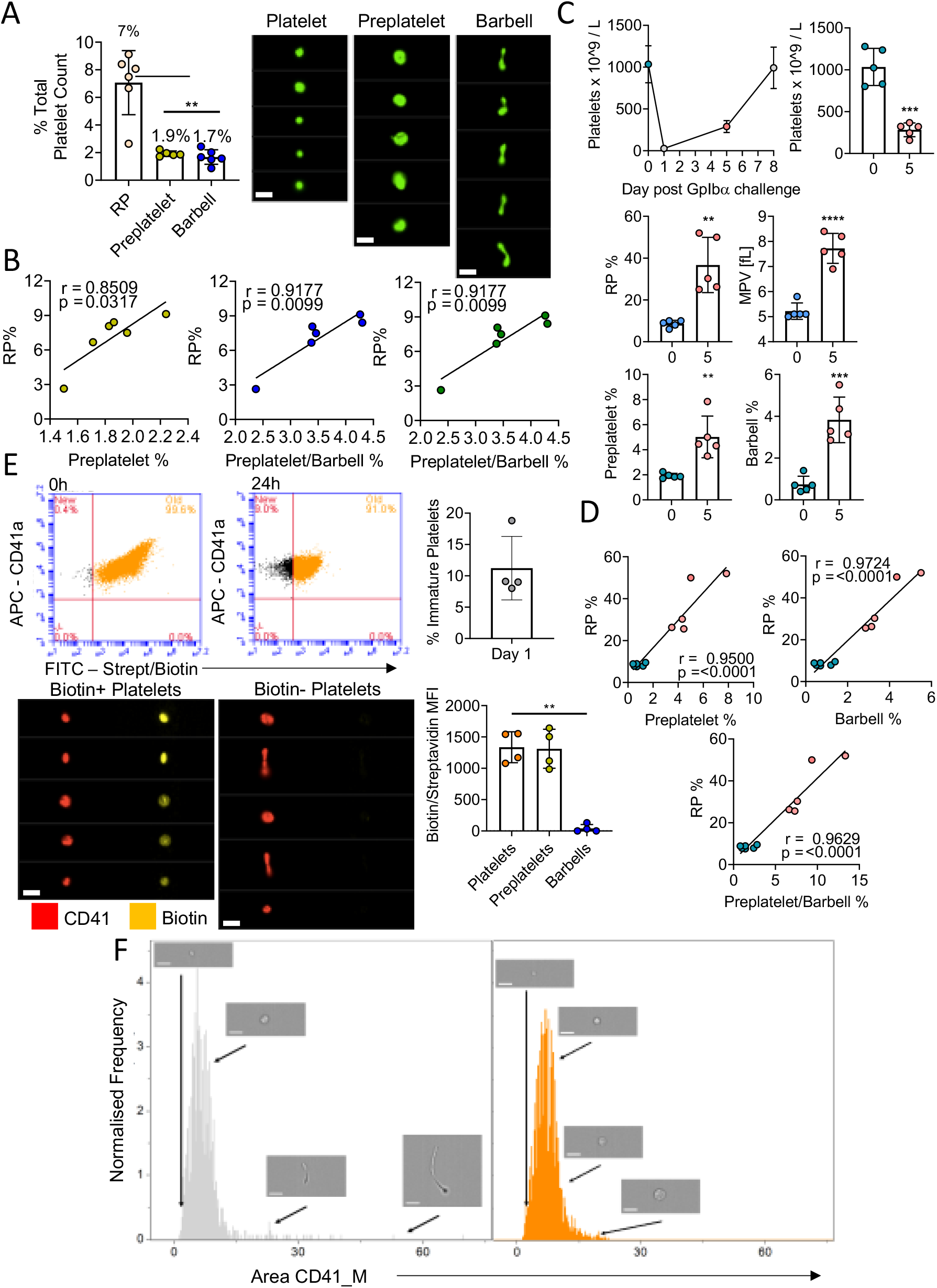
Preplatelets are newly formed immature reticulated platelets. (A)Trisodium citrate whole blood from wild type mice was incubated for 1.5h at 37°C and percentages of reticulated platelets (RP) by flow cytometry, preplatelets and barbells by ISFC using anti-CD61-FITC were quantified. (B) Preplatelets and/or barbells correlated with RP (n=6). (C) Mice (n=5) were treated with a GPIbα polyclonal antibody (1.5μg/mL) and platelet counts and MPV were measured by ABX Pentra 60 (Horiba) haematology analyser and RP by flow cytometry using thiazole orange labelling on day 0, 1, 5 or 7. Also, preplatelets and barbells were quantified by ISFC using FITC CD41 at day 0 (prior to platelet depletion) and day 5 (platelet engraftment) and correlated with reticulated platelets (RP). (Mice (n=4) were injected twice intravenously with NHS biotin (4mg/mL) to label all circulating blood cells with biotin, bled 0 and 24h later and labelled with APC anti-CD41 and FITC conjugated streptavidin. (E) 100% of circulating platelets were verified as biotin positive at baseline and newly formed platelets at 24h post biotinylation were determined as biotin negative. Representative ISFC images show biotin positive and negative platelet morphology (x60 lens, scale bars = 7μm). Biotin mean fluorescence intensity of platelets, preplatelets and barbells was also measured by ISFC at 24h. (F) ISFC to determine the size distribution of biotin negative (immature) and positive (mature) platelets 24h post injection (representative images are depicted with brightfield ISFC imaging, x60 lens, scale bars = 7μm). (A & E) one-way anova with Tukey test; (B & D) Pearson’s correlation coefficient; (C) unpaired t-test. Sig. ** <0.01; ***<0.001; ****<0.0001. +/- 1 SD.

To characterise these structures in greater detail, α-tubulin-labelled PRP was imaged using super-resolution structured illumination microscopy(SIM). Barbell platelets consisted of a marginal band with microtubules extended through the narrow cytoplasmic bridge or shaft and looped back at the distal ends to form two teardrop-shaped structures(Figure 1B, blue arrows). Large preplatelets(yellow arrows) and a new population of intermediate elongated preplatelets, characterised by an oval-shaped marginal band (purple arrows), were resolved which appear to be in the process of forming barbells. Preplatelets and barbells can be further distinguished from platelets by their greater granularity, as measured by ISFC side scatter(SSC), labelling with the immature platelet markers thiazole orange (TO) and HLA-1, and using mitotracker which detects platelets containing greater numbers of mitochondria (Figure 2A). Using the ISFC protocol shown in the Supplementary Figure 3 (with elimination of CD62p positive platelet doublets), the mean number of preplatelets and barbells in whole blood was 3.3+1.6% and 0.5+0.27% (n=12) of the platelet population, respectively, correlating with the %IPF (Figure 2B). These data suggest that preplatelets and barbells are immature.

### Preplatelets and barbells in acquired thrombocytopenia

Preplatelets and barbells were quantified by ISFC within whole blood from patients with ITP(n = 7) relative to controls(n=12). ITP is characterised by low platelet counts(mean 46.5+36.9 × 10^9^/L versus 250.9+52.5 × 10^9^/L), a higher MPV (mean 12.3+0.8 fL versus 9.7+0.8 fL) and higher percentage of newly formed platelets (IPF mean 20.5+12.7% versus 4.1+1.7%). The percentage of preplatelets(10.5+4.7%) and barbells(1.7+1.6%) in ITP was significantly(p<0.05) greater than controls(3.2+1.5% and 0.46+0.21%, respectively), and correlated to the increase in the %IPF (Figure 3A). While preplatelets and barbells in ITP show similar labelling for tubulin(Figure 3B) they were significantly(p< 0.05) larger than in controls(∼10% increase, respectively; Figure 3B and C).

Chemotherapy (in the context of autologous stem cell transplantation) mediates bone marrow ablation and terminates platelet production and can therefore be used to verify whether preplatelets and barbells are part of the newly formed IPF. Preplatelets and barbells were quantified within whole blood from patients (n=6) with lymphoma and myeloma prior to and post chemotherapy. The patients’ platelet count and %IPF prior to bone marrow ablation were 231+27 × 10^3^/μL and 3.2+0.74%, respectively. Preplatelets and barbell levels were 3.8+1.3%, and 0.45+0.04%, respectively(Figure 3D). Post-chemotherapy, the platelet counts were significantly reduced(35.0+35.5 × 10^3^ /μL), but surprisingly the proportion of IPF and preplatelets was not significantly different(2.0+1.1% and 3.3+1.3% respectively). In contrast, barbells could not be detected (0.08%+0.06; Figure 3D) suggesting they are formed from immature platelets which will be absent after bone marrow ablation. Large platelets (>3μm in diameter) must also therefore represent mature platelets and are a heterogeneous population of different ages including immature preplatelets.

### Preplatelets convert into barbells that can undergo fission

Previous studies report that preplatelet and barbell transformation is promoted when incubating washed platelets under cell culture conditions within serum free media at 37°C.^21^ To study barbell formation *in vitro* by ISFC, human control citrate blood and washed platelets were incubated at 37°C in serum-free M199 media for up to 24h. Under these conditions, barbell formation increased approximately 4x in washed platelets compared to whole blood up to 3h and perimeter measurements and tubulin labelling confirm they originate from platelets >3μm in diameter and share a similar morphology as human barbells(Figure 4A and Supplementary Figure 1-3). Figure 4B demonstrates barbells can be detected at 1.5, 3 and 6h but are completely absent by 24h. There was an inverse relationship between decreasing numbers of preplatelets with increasing barbells over time (Figure 4B). Dual labelling of washed platelets with cytosolic CellTrace™ green or red cytosolic dyes confirm that barbells originate from single large platelets(Figure 4C).

To further track the fate of preplatelets and barbells, washed platelets were labelled with CellTrace™ and incubated for 24h at 37°C. Baseline measurements confirm that 100% of platelets were positive for CellTrace™. After 24h incubation, a new population of platelets appeared with significantly reduced CellTrace™ labelling(Figure 5A). The preplatelets appeared to undergo at least two rounds of fission to form 11% more platelets, coupled with a decrease in platelet size. Newly formed platelets not only exhibited higher Mitotracker labelling(Figure 5B) but the mitochondrial electron transport chain inhibitor rotenone(3μg/mL) inhibited barbell formation, suggesting mitochondrial function and energy is important(Figure 5B). Furthermore, barbell formation was also inhibited by nocodazole and cytochalasin D(Figure 5C). Preplatelets are therefore capable of undergoing fission, which is dependent on cytoskeletal remodelling and mitochondrial respiration.

### Preplatelets and barbells in murine blood

To confirm that preplatelets and barbells are not exclusive to human blood, we performed measurements in mouse blood and anti-platelet antibody-treated blood as a model of ITP. We used *in vivo* biotinylation labelling to conclusively confirm whether preplatelets and barbells represent newly formed immature platelets. Using a similar ISFC protocol(Supplementary Figure 4A), preplatelets and barbells (with similar tubulin cytoskeletal structure and morphology to human) were also detected in murine blood under the same conditions(Supplementary Figure 4B-D) such as temperature and anticoagulant as humans(1.9%+0.18 and 1.7%+0.48 respectively) and correlate with immature RP levels(7%+2.1)(Figures 6A and 6B). WT mice were treated with anti-GPIbα to induce severe thrombocytopenia and the time course of recovery of platelets was monitored, to investigate the temporal relationship between preplatelets, barbells and platelets. Figure 6C shows the kinetics of the platelet count with a nadir at day 1, increasing to 288 × 10^3^ /μL at day 5 and near normal by day 7. At day 5, the percentage of RP and MPV increased by 76.5% and 32.5%, respectively) confirming the platelet population consisted predominantly of immature platelets(Figure 6C). Preplatelets and barbells were not only larger than baseline(Supplementary Figure 5) but also significantly increased in number and correlated with the fraction of RP at days 0 and 5(Figure 6D). To conclusively confirm whether preplatelets and barbells are immature newly formed platelets, mice were intravenously injected twice with NHS biotin to ensure that 100% of circulating platelets were labelled (Figure 6E). At 24h 11.2%+5.1 of circulating platelets were shown to be biotin negative and thus represent newly formed immature platelets (Figure 6E). The barbell structures were all biotin negative confirming that they originate from immature cells. In contrast, preplatelet(>3μm in diameter) and other circulating platelets (<3μm in diameter) were positive for biotin(Figure 6E-F). Moreover, the biotin-negative platelets were not exclusively large, confirming their heterogeneity(Figure 6F).

Preplatelets and barbell measurements in both human and murine normal and thrombocytopenic blood correlate with measurements of platelet immaturity. Furthermore, *in vivo* studies in mice confirm that barbells are exclusively derived from a population of immature large preplatelets, however, not all large platelets are immature.

## Discussion

Despite the accumulating evidence that platelets can undergo fission,^21,23^ it remained unclear whether all circulating platelets were capable of division and if the number of the precursors changed in conditions of increased or decreased platelet production. We hypothesized that immature RP are equivalent to preplatelets and can undergo fission. In this study we have therefore utilised the power of ISFC and microscopy to both characterise and quantify preplatelet maturation and to measure immature platelets both *in vivo* and within *ex vivo* cell culture. We confirmed barbells were indeed present in citrated whole blood(Figure 1) and exhibit a continuous figure-of-eight tubulin cytoskeleton with two distal teardrop-like structures interconnected by a cytoplasmic bridge. Interestingly, barbells were present in human blood immediately after phlebotomy(Figures 1 & 2) and their numbers remained constant for 3h(Figure 4A) suggesting that they could be undergoing continuous maturation and division. To determine if preplatelets can undergo barbell formation and eventual fission *in vitro*, human control washed platelets were incubated under cell culture conditions. Barbell formation was observed up to 6h, but was absent by 24h. Barbell conversion was inhibited by both nocodazole and cytochalasin D confirming the importance of the cytoskeleton, namely tubulin and actin, in platelet maturation(Figure 4B).^23^ Dual labelling experiments with cytosolic CellTrace™ green or red dyes also confirmed that the barbells originated from single platelets(Figure 4C). Furthermore, cell tracking experiments confirmed the appearance of a new population of platelets with reduced fluorescence(Figure 5A). Remarkably, preplatelets could also undergo at least two rounds of fission signified by the two separate peaks to result in 11% more platelets.

We then clearly identified both preplatelets and barbells within normal human and mouse whole blood. Furthermore, these structures represent comparative subpopulations(3.6-3.8%) of the total platelet count, mirroring IPF/RP measurements and expressing immature platelet markers HLA-I and high levels of thiazole orange fluorescence. The overall percentage of barbell structures in human whole blood was low(∼0.5%), suggesting that preplatelet maturation is transient and likely to be enhanced under normal shear and turbulent forces within the vasculature. ^17,23,25,26^ Despite this, the fraction of barbells observed in our study in human whole blood was still ten-fold greater than previous observations,^23^ using labelled PRP on slides using a laser scanning cytometry approach. This difference possibly reflects the differences between ISFC and microscopy in detecting rare, transient, and fragile structures.^27-29^ Furthermore, the frequency of preplatelets and barbells also correlated with the IPF and RP measurements suggesting that this is also an indirect measure of the rate of platelet production both in controls and acquired thrombocytopenia. Indeed, the percentage of preplatelets and barbells in ITP was significantly greater than controls (Figure 3A), but *ex vivo* barbell formation also correlated with the %IPF. Tubulin labelling confirmed that the frequency and size of preplatelets and barbell structures in ITP was much greater than with healthy controls (Figure 3B).

Preplatelets and barbells were validated as immature platelets in a variety of ways. As expected, *in vivo* antibody platelet depletion experiments demonstrated an increase in RP and MPV at day 5 post-injection of anti-GPIb(Figure 6C) when platelet production was maximal a few days before the normal count was restored. ISFC not only confirmed that the preplatelets and barbell structures were also significantly increased but that the percentages correlated with the fraction of RP at days 0 and 5(Figure 6B).

*In vivo* biotinylation labelling of the circulating pool of platelets in mice was performed to monitor the kinetics of ageing and immature platelet production. This method was previously used to confirm that RP as measured by CFC are indeed the youngest platelets.^3,30^ Analysis of post biotin labelling blood at 0h confirmed that all circulating platelets were indeed biotin positive(Figure 6E). However, at 24h post biotinylation, ISFC confirmed that all circulating barbells were completely biotin negative.

The IPF provides an indirect measurement of the rate of thrombopoiesis and high or low IPF values therefore signify either enhanced bone marrow activity or suppression respectively. ^8,9,31^ However, IPF often fails to distinguish between healthy individuals and those with conditions of bone marrow suppression/failure.^10,32-35^ Moreover, analysis of various macrothrombocytopenias often gives very high IPF values, probably caused by non-specific labelling of the large platelets, and is therefore unrelated to the rate of thrombopoiesis.^11^ In this study, within whole blood from ITP patients we observed that the fraction of preplatelets and barbells correlated with increased platelet turnover and importantly, in response to bone marrow ablation, barbell formation was absent. Interestingly both the percentage of preplatelets and IPF(platelets 3-10μm in diameter) were not significantly reduced following bone marrow ablation. Furthermore, our biotinylation experiments not only demonstrate that not all platelets >3μm in diameter are immature but importantly that newly formed platelets can also vary in size(Figure 6). Therefore, not all large platelets have the capacity to transform into barbells and duplicate. Although platelet size is an important characteristic of thrombocytopenia with peripheral destruction(e.g. ITP), it is historically reported in the literature under resting conditions that platelet size and age do not necessarily correlate. ^36,37^

For preplatelets to convert into barbells they must overcome marginal band elastic bending forces and cortical actin-myosin compression. As the marginal band consists of on average 8-10 microtubule bundles, larger platelets have a thinner marginal band to overcome such force constraints.^23,38^ In agreement with this filamin A KO mice characterised by a thicker marginal band exhibit a defect in preplatelet/barbell formation. Such findings have left many questions open regarding which macrothrombocytopenias may have a defect in platelet maturation and morphogenesis.^39^ ITP is an acquired macrothrombocytopenia and in subjects with α_IIb_β_3_ autoantibodies, MK proplatelet formation is suppressed yielding fewer and denser extensions compared to normal.^40,41^ However, as we observed an enhancement in preplatelet and barbell formation in ITP patients this would suggest platelet marginal band morphology is normal and such exacerbations in barbell formation are due to increases in MPV. ^42^

Considering all the findings in this paper, an updated model of preplatelet maturation within the circulation is illustrated in Figure 7. We demonstrate preplatelet maturation occurs only in newly formed immature platelets and the rate in which this takes place signifies the rate of platelet turnover in health and disease. In addition to platelet counts and IPF, morphometric quantification of preplatelet-derived barbells may provide an additional tool for assessing the cause of thrombocytopenia and the individual’s capacity to recover their low platelet count. By using appropriate anticoagulants and labelling, image analysis of large numbers of platelets on blood films or by ISFC could be used to measure barbell structures in whole blood.

**Figure 7:**
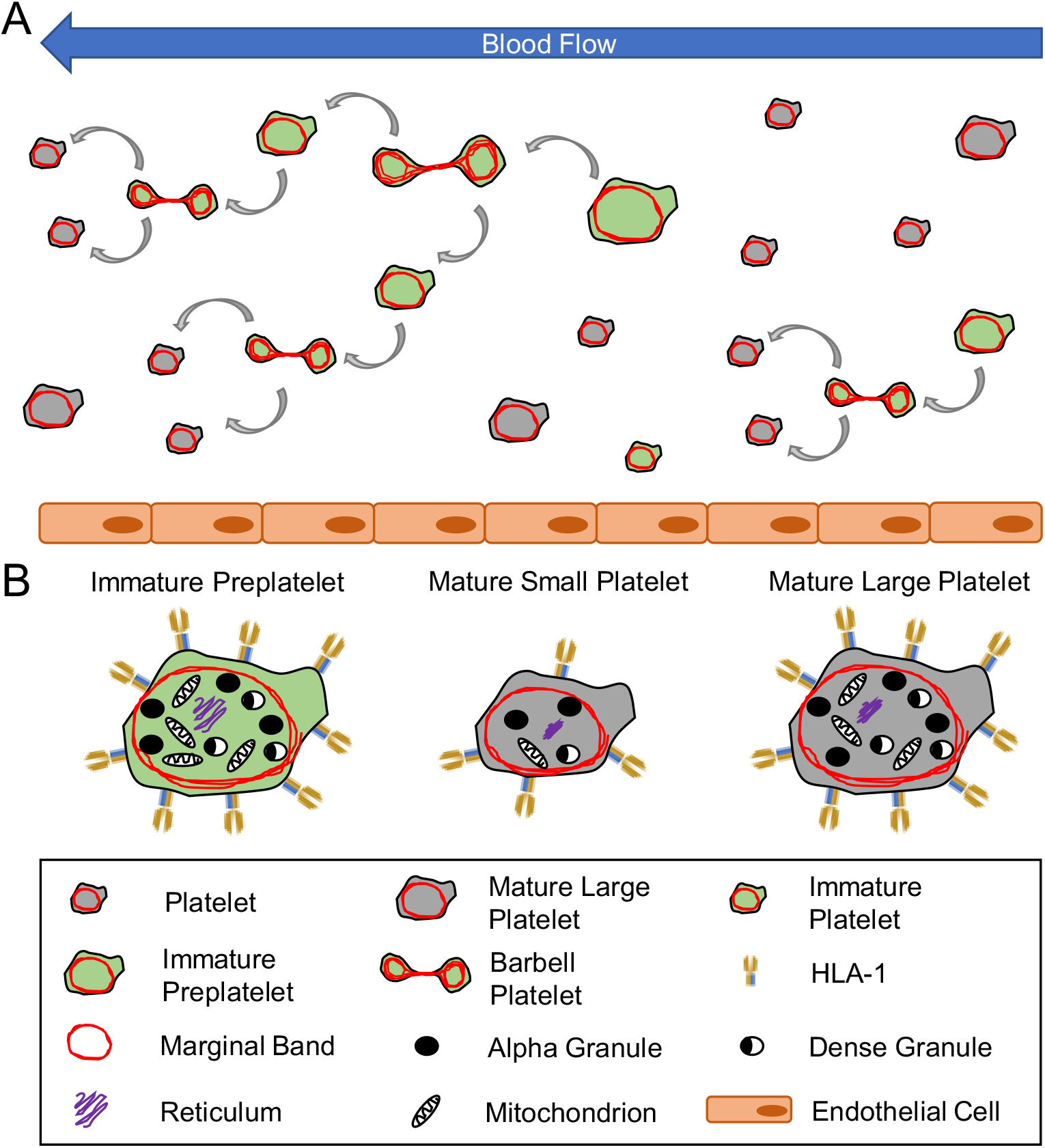
Model of Platelet Maturation in the Bloodstream. In (A) circulating platelets are heterogeneous in size and age. Immature platelets >3 μm in diameter are termed preplatelets (determined by a green cytoplasm). These platelet progenitor cells mature by continuously transforming into barbell platelets and undergoing fission into two smaller platelets until reaching a size threshold of <3μm in diameter. Under steady state production, not all large platelets are immature. Unlike preplatelets, these lack the capacity to undergo maturation. Therefore, mature platelets consist of small and large platelets (determined by a grey cytoplasm). (B) Barbell platelets originate from immature preplatelets consisting of a greater nucleic acid content, granule content, number of mitochondria and HLA I expression compared to mature small platelets. In contrast, mature large platelets contain similar numbers of granules and mitochondria which are non-specifically labelled with dyes used for measuring IPF and RP. Large mature platelet express slightly less HLA I than immature preplatelets which could potential separate preplatelets from mature large platelets.

## Acknowledgements

This project was funded by a British Heart Foundation 3-year PhD studentship grant (FS/17/29/32828) and supported by a BHF Accelerator Award (AA/18/2/34218) and by the CRN (Clinical Research Network West Midlands). We thank Dr. Jun Mori and Dr. Timo Voegtle for their advice and technical help with the mice studies. We also thank Dr. Charles Percy, Dr. Hayder Hussein, Dr. Will Lester, Dr. Suzanne Morton and Beth Lovell for their clinical input and help with patient recruitment. The authors would like to acknowledge the Imaging Suite at the University of Birmingham for support of imaging experiments. Imaging facilities used in this project were funded by the University of Birmingham, COMPARE and the BHF.

## Authorship Contributions

SK and AD generated the data and wrote the paper. PH conceived, funded the project and wrote the paper. GL and PN provided clinical input and clinical samples for the study and edited the paper. ST and SPW provided advice and edited the paper. YS provided mice edited the paper and contributed to funding.

