## Supplementary for "Analysis of Preplatelets and Their Barbell Platelet Derivatives by Imaging Flow Cytometry"

**Supplementary Methods**

***Patient recruitment and demographics***

Healthy volunteers contained varied UK ethnic backgrounds with no known healthy problems and were aged between 18 and 60. Patients with primary, chronic ITP donated blood on a one-time basis only and consisted of males and females with varied UK ethnicities, an age range between 18 and 80 and were treatment naïve (n=3) or varied daily or weekly doses of prednisone and/or a TPO mimetic (romiplostim or eltrombopag). In contrast, patients with myeloma or lymphoma (with varied UK ethnicity, aged 41 – 68) were receiving chemotherapy which consisted of a single high dose of melphalan or six days of LEAM (**L**omustine, **E**toposide, cyt**A**rabine and low dose **M**elphalan) followed by an autograft stem cell transplantation on the subsequent day post therapy. Blood samples were taken before chemotherapy (i.e. at baseline) and 6 days following the autograft transplant (i.e. at the platelet count nadir: with no platelet production).

Patients were excluded from the study if thrombocytopenia was caused by an alternative condition to ITP, patients had a suspected congenital disorder, received a blood transfusion within 7 days prior to recruitment, were currently receiving treatment known to affect platelet function and/or diagnosed with severe anaemia.

***Flow cytometry***

All Samples were diluted to a final concentration of 1/1000 prior to Accuri^TM^ C6 acquisition. To determine reticulated/immature platelets manually by flow cytometry, 5μL of murine whole blood was labelled with anti-mouse CD41 APC for 15 min. Post incubation, BD Retic-count (TO) was added at a concentration of 10μg/mL and incubated for 30 min. Samples were fixed in 2% formalin. Reticulated platelets were determined by forward scatter and TO fluorescence intensity. To determine biotin positive platelets by flow cytometry, 5μL of whole blood was labelled with anti-mouse CD41 APC and FITC streptavidin for 15 minutes at room temperature and fixed with 2% formalin in PBS. To define biotin/streptavidin positive platelets, gates were generated based on a negative control (NHS-biotin naïve mice).

***Immunofluorescence microscopy***

PRP was incubated under the same conditions as above for 1.5h, fixed in 2% formalin in PBS, diluted to 5 x 10^6^ and centrifuged at 200g for 10 minutes onto poly-L-lysine (0.01%; Sigma, A-005-C) coated 13mm coverslips (VWR; 631-1578P). Platelets were permeabilized in 0.1% Triton X-100 (Sigma, T8787) for 10 minutes, and non-specific binding was blocked with 1% goat serum (Sigma, G9023) and 2% BSA (Sigma, A9647) in PBS for 1 hour. Tubulin was labelled with an anti-mouse-tubulin monoclonal antibody (1/500; Sigma, T8203) in 2% BSA, 1% goat serum in PBS for 1 hour to avoid species cross absorption and secondary labelled with goat anti-mouse IgG (H+L) alexa fluor 488 (1/1000; Invitrogen, A-11001) for 30 minutes. Between permeabilization, blocking and antibody incubation steps, platelets were washed three times with PBS. Following secondary labelling, coverslips were mounted in Vectashield vibrance aqueous solution (Vector, H-1700) and platelets were imaged by widefield using either Leica DM6000 or Structured Illumination Microscopy (SIM).

For SIM imaging, samples were imaged and processed on a Nikon N-SIM-S microscope with Ti2-E inverted microscope stand, PFS4 focus stability system, 100x 1.49 NA oil immersion objective lens with automatic correction collar adjustment, LU-NV-L laser bed, Hammamatsu Orca Flash 4 V3 sCMOS camera, TI2-FT N-SIM Motorized Filter set, and NIS-AR elements V5 software. For Leica DM6000 B microscopy, widefield imaging with 60x1.6 objective lens, Leica application suite with DMC control for windows^TM^ 2000 XP.

**Supplementary Data**

**Results**

***Optimisation of Pre-analytical variables for measuring preplatelets and Barbells***

To determine the optimal conditions for measuring preplatelets and barbells, human whole blood (n=5) in different anticoagulants was immediately (post phlebotomy) incubated with either CD61 for 15 min at 21 or 37°C or for live tubulin labelling CD61 and AF674 SiR tubulin (4μM) for 30 min. Platelet morphology was manually interrogated by plotting aspect ratio (a measurement of circularity) against area of CD61 fluorescence using ISFC. Platelets and preplatelets were present in all anticoagulants at both temperatures (Supplementary Figure 1A, Images 1 & 2). Platelets in EDTA blood remained compact and discoid with no barbell-shaped objects. Barbells (two distal platelet interconnected by a narrow cytoplasmic bridge) were only present in 37°C citrate (Supplementary Figure 1A, Image 4). The barbell cytoskeletal structure is described in detail in Figure 1. At both temperatures, citrate (and Hirudin) blood also contained elongated barbell-shaped platelet microaggregate contaminates which appeared to consist of two separate marginal bands (Supplementary Figure 1A Image 3 and Supplementary Figure 1B). Blood smears labelled under the same conditions as above also support these findings (Supplementary Figure 1C, red arrows: microaggregates & blue arrows: barbells), identifying 37°C citrate the optimal conditions for characterising and quantifying preplatelets and barbells in blood.

To determine if microaggregates are indeed barbells undergoing a different transitional of preplatelet/barbell transformation or merely barbell-shaped contaminates, citrate blood was labelled with CD61 and CD62p and microaggregates were quantified at 21°C up to 3h post phlebotomy. Microaggregates (as did platelet activation) significantly increased with time (Supplementary Figure 2A). CD62p labelling demonstrated these structures consist of at least one activated platelet (Supplementary Figure 2B). The concentration of microaggregates also positively correlated with CD62p MFI (Supplementary Figure 2C) and were absent in citrate blood containing 10μM eptifibatide (an inhibitor of α_IIb_β_3_-mediated platelet aggregation; Supplementary Figure 2D). to determine CD62p exposure on barbells, citrate whole blood was incubated at 37C for 15 min. Barbells were CD62p negative Supplementary Figure 2E). Interestingly, barbells were also absent (Supplementary Figure 2F). As there was no difference observed in platelet CD61 MFI with or without eptifibatide this confirmed eptifibatide has a direct inhibitory effect on barbell formation rather than interfering with the CD61 fluorescent signal (particularly at weaker regions of barbells such as the narrow cytoplasmic bridge). In summary, citrate blood contains barbells and pseudo-activated barbell-shaped microaggregates which can be discriminated from true barbell by CD62p expression.

From the above findings an ISFC gating strategy was designed to further characterise and accurately quantify preplatelets and barbells in 37°C citrate whole blood. Both preplatelet and barbell gating was based on 10,000 CD61 positive platelet images and microaggregate contaminates were excluded by gating out CD62p positive events (Supplementary Figure 3A). For preplatelets, single, circular CD61 focussed / CD62p negative platelet images were initially discriminated using aspect ratio (0.8-1) and area of a CD61 morphology mask generating the “Circular” gate. From this, using the CD61 erode+4 mask, preplatelets were determined by a diameter of 3-10μm. For barbells, CD61 positive / CD62p negative elongated platelets images were separated from all other platelet images plotting the feature compactness against combined features symmetry2 multiplied by height (Sym2*H). Barbell platelets were subsequently separated from the “elongated” population using minor axis intensity of CD61 morphology against the area of CD61 morphology multiplied by the major axis intensity (Min-Ax-Int M) of a CD61 skeleton thin mask (A*Maj-Ax-Int STn). Supplementary Figure 3B shows barbell-shaped microaggregates are removed from the barbell gate with CD62p gating.

Using the above ISFC gating strategy, morphometric analyses show compared to platelets, the mean preplatelet diameter is significantly larger at ~3.4μm, with a perimeter range of 13-24μm compared to (Figure 1C). Thus, 13μm perimeter measured by ISFC equates to the minimum 3μm preplatelet diameter. Interestingly, the barbell perimeter was 13-42μm with a significantly greater mean perimeter than preplatelets. Together these data not only confirm the larger preplatelets have already transformed into barbells but the smallest barbell perimeter (13μm) equated to that of preplatelets agreeing with others that barbells originate from preplatelets $\geq$3μm in diameter.^23^

**Supplementary Figure 1: Platelet morphology is dependent on anticoagulant and temperature.**

(A) Human control whole blood from the same donor was anticoagulated in EDTA, Hirudin and trisodium citrate and labelled with FITC ant-CD61 for 15 min or FITC anti-CD61 and SiR tubulin (4μM) for 30 min at 21 and 37°C. 10,000 CD61+ platelet images were acquired using ISFC and platelet morphology was depicted by aspect ratio and area of a CD61 fluorescence morphology. Images of platelets, preplatelets, barbell-shaped microaggregates and barbell platelets are depicted using CD61 and SiR tubulin fluorescence (n=5; scale bar = 7μm, x60 lens). (B) SIM microscopy of tubulin labelling of microaggregates (red arrows) from citrate anticoagulated PRP prepared at 21°C (x100 magnification, n=3, scale bar = 5μm). (C) Blood from the same healthy donor was anticoagulated in EDTA, hirudin and 10% trisodium citrate and maintained at either 21 or 37°C whilst preparing for blood smears (n=3; x60 lens, scale bar = 5μm).

**Supplementary Figure 2: Barbell-shaped platelet microaggregates are an artefact of platelet pseudoactivation**

Citrate and EDTA blood from the same healthy control donor was labelled with FITC CD61 and BV421 CD62p and incubated at 21°C for 3h, 10,000 CD61 positive platelets images were acquired by ISFC. (A) Concentration of microaggregates and change in platelet CD62p exposure over time(n=5). (B) ISFC images of microaggregates depicting location of CD62p exposure. (C) Correlation between concentration of platelet microaggregates and CD62p MFI (n=5). (D) Citrate whole blood was withdrawn with or without eptifibatide (10μM), incubated at 21^°^C for 3 h and microaggregates quantified by ISFC at 0, 1 and 3h (n= 5). (E) Healthy donor citrate blood was incubated immediately with FITC anti-CD61 and BV421 anti-CD62p for 15 min at 37°C and barbells were image using ISFC to determine CD62p exposure. (F) under the same conditions, citrate whole blood was withdrawn with or without eptifibatide (10μM) to determine the presence of barbell platelets by ISFC and total platelet CD61 MFI (n=5). (A & D) Two-way anova with Bonferroni multiple comparisons test, (A & C) Pearson’s Coefficient; (F) paired t-test Sig. *<0.05, **<0.01, ****<0.0001. +/- 1 SD.

**Supplementary Figure 3: Using ImageStream MKII to identify preplatelets and barbells**

Healthy control citrate whole blood was incubated immediately with FITC anti-CD61 and BV421 anti-CD62p for 15 min at 37°C. (A) CD62p positive platelets (including barbell-shaped microaggregates) were excluded from the analysis. For preplatelets (left side gating): circular CD61 platelet images were gated by aspect ratio (0.8 – 1) and area of a morphology (M) mask. Platelets (light blue; <3μm in diameter) and preplatelets (yellow; >3μm in diameter) were discriminated and quantified from the “Circular Platelets” population using an erode+4 mask (E4) with the feature diameter. For barbell platelets (right side): elongated platelet structures were initially separated from single circular or discoid platelet images using compactness and symmtery2 multiplied by height (Sym2*H) of CD61 morphology. Elongated platelets were further interrogated by minor axis intensity (Min-Ax-Int) CD61 morphology against area multiplied by major axis intensity (A*Maj-Ax-Int) CD61 skeleton thin (STn) generating the “barbells” region. Also, ISFC images (x60 lens) of platelets, preplatelets and barbell platelets using brightfield (BF) and CD61 fluorescence. (B) In the absence of BV421 CD62p gating, barbell-shaped microaggregates are present within the barbell gate (red dots) in citrate blood at 37°C. As these microaggregates express CD62p (see images), they can be removed from the barbell population by gating out CD62p positive events. Scale bar 7μm. (C) Diameter and perimeter measurements of preplatelets and barbells quantified by IFC using a CD61 erode+4-pixel mask. Freq: frequency. (D) Paired t-test. Sig ****=<0.0001. +/- 1 SD.

**Supplementary Figure 4: Characterisation of murine preplatelet and barbells**

(A) Preplatelets and barbell platelets were discriminated using the same gating strategy described in Supplementary Figure 3. (B) Murine blood was anticoagulated with EDTA or trisodium citrate (n = 6), incubated for 1.5h at 37°C, and labelled with BV421 anti-CD62p, FITC anti-CD41a and/or SiR tubulin (4μM). 10,000 CD41a platelet images were acquired using ISFC (n=6). (C) representative ISFC images of platelets, preplatelets and barbells depicted by CD41a and SiR tubulin. (D) Diameter of platelets and preplatelets and perimeter measurements of preplatelets and barbells quantified by IFC using a CD41 erode+4-pixel mask. E4: erode+4, Min-Ax-Int: minor axis intensity, M: morphology, A*Maj-Ax-Int: area multiplied by major axis intensity, STn: skeleton thin. (D) Paired t-test. Sig ****=<0.0001. +/- 1 SD.

**Supplementary Figure 5: Preplatelets and barbells increase in size in response to GPIb-mediated thrombocytopenia**

(A) Mice (n=5) were inflicted with GPIbα-mediated platelet destruction and preplatelet and barbell mean area (μm^2^) was determined at day 0 (prior depletion) and 5 (platelet engraftment) post platelet depletion by IFC using a erode+4pixel mask of CD41 fluorescence (representative images of preplatelets and barbells are depicted by FITC anti-CD41 fluorescence, x60 lens, scale bars = 7μm). Unpaired t-test. Sig ***=<0.001. +/- 1 SD.

Supplementary Figure 1

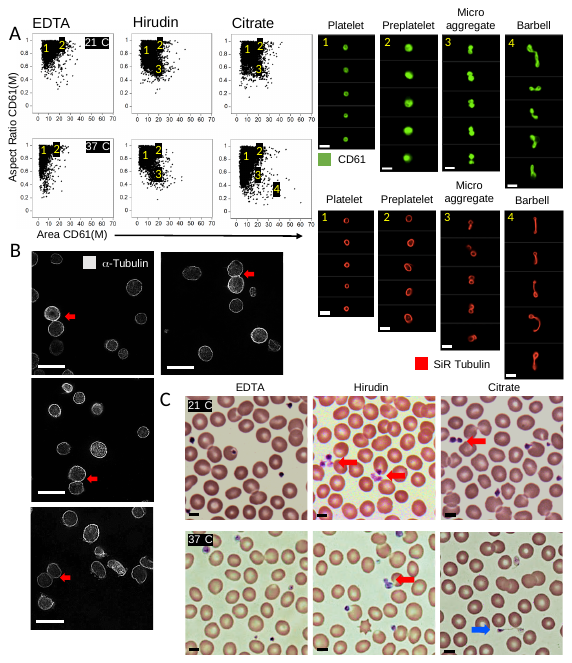

Supplementary Figure 2

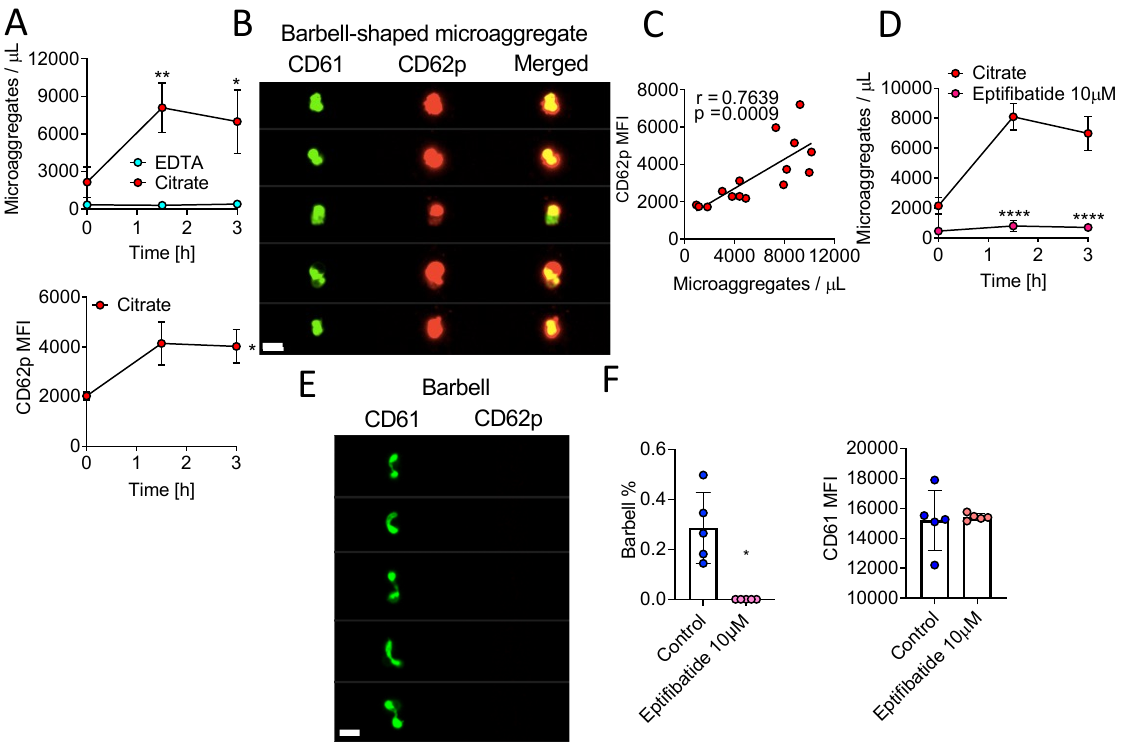

Supplementary Figure 3

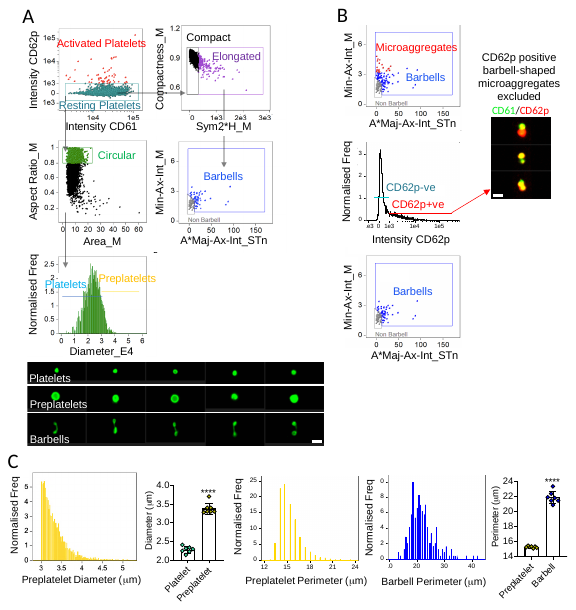

Supplementary Figure 4

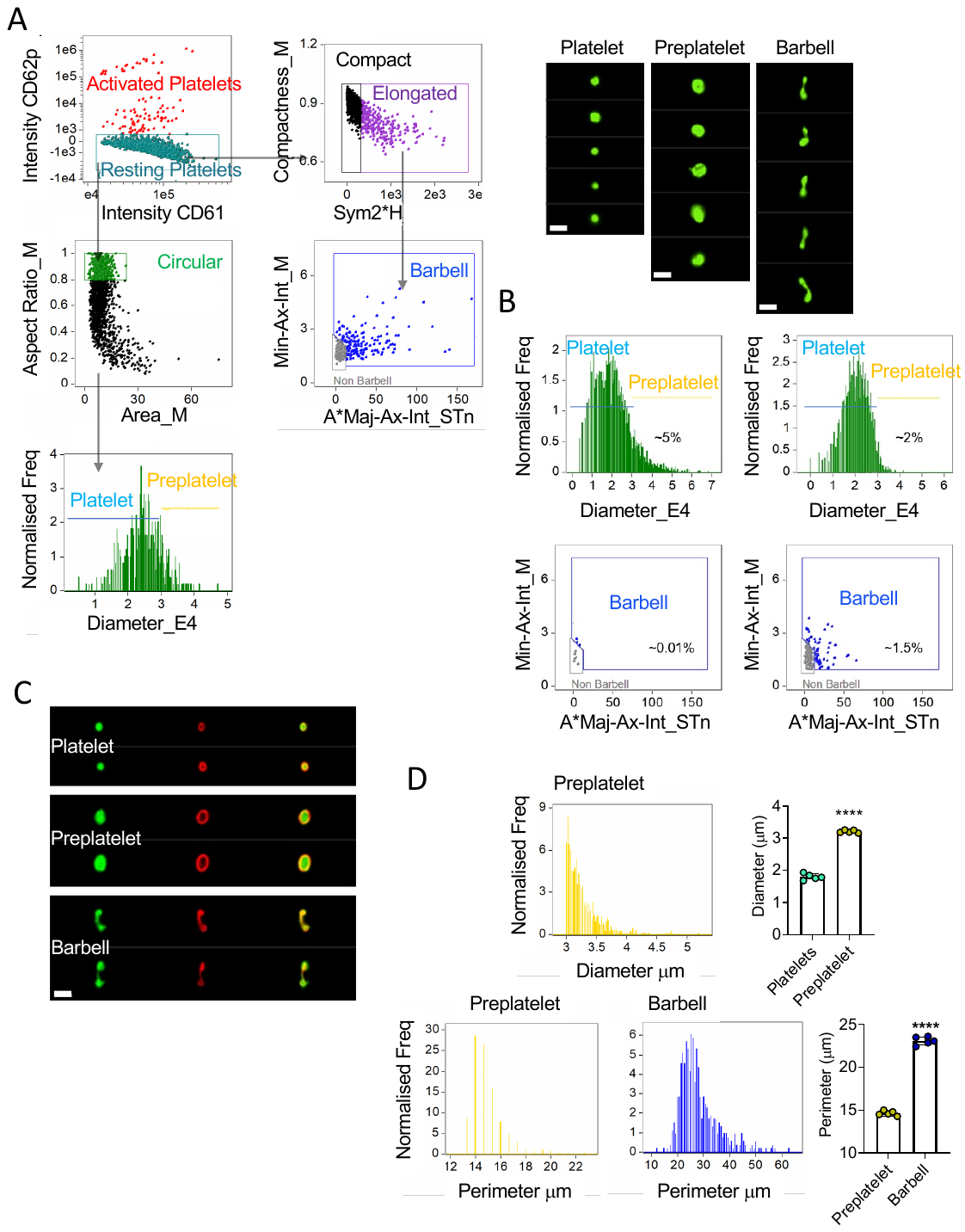

Supplementary Figure 5

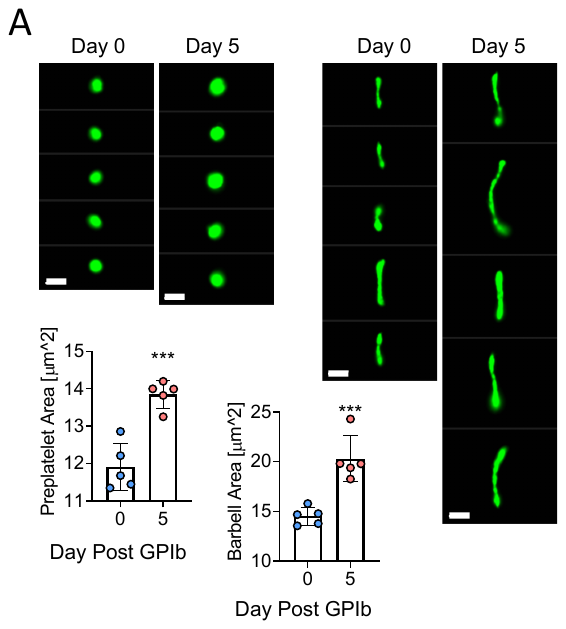
